# Nanopore sequencing reaches amplicon sequence variant (ASV) resolution

**DOI:** 10.64898/2026.02.26.708165

**Authors:** Marie Riisgaard-Jensen, Stefania Andrea Rosso Villanelo, Kasper Skytte Andersen, Rasmus Kirkegaard, Susan Hove Hansen, Chenjing Jiang, Albert Vejlin Stefansen, Jakob Holm Dalsgaard Thomsen, Per Halkjær Nielsen, Morten Kam Dahl Dueholm

## Abstract

Sequencing of ribosomal marker genes remains a cornerstone for profiling complex microbial communities. In recent years, there has been a shift from Illumina to long-read technologies, including PacBio and Oxford Nanopore Technologies (ONT). ONT is attractive due to its low startup cost and portability; however, historically high error rates have prevented direct amplicon sequencing variant (ASV) generation from raw nanopore reads. This has forced most workflows to rely on mapping raw reads against reference databases constraining analyses to taxa covered by these. With recent improvements in ONT sequencing accuracy, we sought to challenge this view by sequencing samples of increasing complexity using primer sets targeting amplicons of different lengths, and by sequencing the exact same PCR libraries on both PacBio and ONT. We demonstrate that error-free ASVs can now be generated directly from raw nanopore reads using standard denoising algorithms originally developed for Illumina data. Current ONT read quality enables reliable reconstruction of amplicons spanning ∼250 bp to ∼4,200 bp and allows resolution of intragenomic rRNA gene variants. These results extend beyond simple mock communities to complex fecal, anaerobic digester, activated sludge, and soil samples. When sequencing depth is sufficient, ONT accurately recovers all or nearly all intra-genomic 16S rRNA gene copy variants, showing perfect sequence identity to curated reference sequences in mock communities and to ASVs inferred from PacBio data in complex communities. Across the primer sets, ONT required higher sequencing depth than PacBio to fully resolve the communities, with this requirement increasing with amplicon length. For complex samples, ONT required approximately 2-3× more reads for V4 (∼250 bp) and V1-V3 (∼500 bp), 4.1-5.6× more reads for V1-V8 (∼1400 bp), and 25-42× more reads for rRNA operon (OPR) amplicons (∼4200 bp). Consequently, sequencing complex communities with OPR primers on ONT is currently not feasible due to the unrealistically high read depth required. This study provides evidence that ONT amplicon sequencing has matured to the point where true ASV-resolved profiling is practically and economically feasible, moving ONT amplicon analysis beyond reliance on OTU clustering or reference alignment to enable application in environments lacking comprehensive reference databases.

**Key Findings:**

1. It is now straightforward to generate ASVs on ONT platforms (250-4200 bp)
2. ONT can resolve intragenomic 16S rRNA gene variants
3. ASV recovery is successful in both simple and complex communities

## Introduction

Amplicon sequencing of the small subunit ribosomal RNA genes is a cornerstone for microbial community profiling across diverse environments. Short-read Illumina sequencing has long been the predominant platform due to its high accuracy and reproducibility. However, the short-read length limits phylogenetic resolution and often prevents confident species identification, which can mask microdiversity in complex communities [1, 2]. Long-read technologies, including PacBio and Oxford Nanopore Technologies (ONT), are increasingly used in microbial ecology as they provide full-length rRNA gene coverage and improved species-level recovery [2]. PacBio offers error rates comparable with short-read platforms [3] while the ONT platform is often chosen for its flexibility, including the ability to sequence native DNA, produce reads exceeding 10 kb, and operate on affordable, portable instruments [4].

Post-processing of amplicon sequences has traditionally relied on either clustering reads into operational taxonomic units (OTUs), usually at 97% sequence identity, or using denoising algorithms to infer exact amplicon sequence variants (ASVs), commonly implemented through tools such as USEARCH and DADA2 [5, 6]. Compared to OTUs, ASVs are generally preferred because they resolve variation at the single-nucleotide level and can be used as consistent identifiers for microbial profiling without relying on a 16S rRNA gene reference database [7]. ASVs have become the standard in Illumina and PacBio workflows and allow for full-length 16S rRNA gene sequences with single-nucleotide resolution [5, 8, 9]. However, the high error rate of ONT nanopore reads has historically made ASV generation practically unfeasible.

Early ONT amplicon workflows were designed to compensate for the higher error-rate by mapping the raw reads against a full-length 16S rRNA gene reference database rather than generating new OTUs or ASVs, making it impossible to detect novel diversity within a sample. As a result, mapping-based approaches rely heavily on the completeness of existing reference databases and prioritize taxonomic classification over variant-level resolution. This strategy is applied by widely used tools such as NanoCLUST and EMU [10, 11]. To accommodate sequencing inaccuracies, these workflows typically apply relaxed identity thresholds, which often limits classification to the genus level despite the availability of long amplicons that could, in principle, support species-level resolution.

More recently, *de novo* clustering approaches such as ONT-AmpSeq, amplicon_sorter, Pike, and CONCOMPRA have been developed [12–15]. These methods use alternative strategies, including OTU generation or UMAP-based clustering, to generate a consensus sequence. Similar to mapping-based approaches, clustering is often done at <98.7% sequence identity, limiting species-level resolution. Additionally, some of these tools rely on extensive read polishing which is computationally intensive and error-prone. Other ONT workflows employ unique molecular identifiers (UMIs) to generate multiple reads of the same original molecule and using the consensus sequence of these to improve accuracy. This technique has been shown to produce error-free *de novo* sequences; however, the increased accuracy comes at the cost of substantially more labor-intensive laboratory protocols and greatly reduced sequencing yield [16–18].

Improvements in ONT, including the continuously improving basecalling models and the introduction of R10.4 flow cell chemistry, have substantially increased per-base accuracy, approaching that of PacBio reads [19–21]. We hypothesize that the recent improvements on the ONT platform enable *de novo* ASV-resolved profiling. This is evaluated by benchmarking ASVs derived from the ONT platform against the PacBio platform using both mock and natural microbial communities of increasing complexity. We further examine how amplicon length and sequencing depth influence ASV recovery, community structure, and comparability across sequencing platforms.

## Results and Discussion

### Complete recovery of expected ASVs in the ZymoBIOMICS mock community

We first evaluated the ASVs generated from both PacBio and ONT sequencing of the ZymoBIOMICS mock community (Zmock) (Cat. No. D6305, Zymo Research, USA), which contains eight bacteria and two fungal species (**Supplementary Table S1**). Only the bacterial component was considered in this study due to primer specificity. For each sequencing platform, we assessed the pipeline across multiple subsampling depths, and the resulting ASVs were mapped to the curated Zmock reference sequences by Lin *et al* (2024) [18] (**Figure 2**). Across all primer sets, exact matches were recovered for all expected theoretical sequences, except for one *Salmonella* V4 variant missing from the ONT dataset. *Salmonella* was the only taxon with two V4 variants; the missing variant differs only by a single base and has a lower copy number (1 vs. 6), making it difficult to resolve with a short amplicon and likely prone to being masked by sequencing errors from the higher-abundance variant.

**Figure 1:**
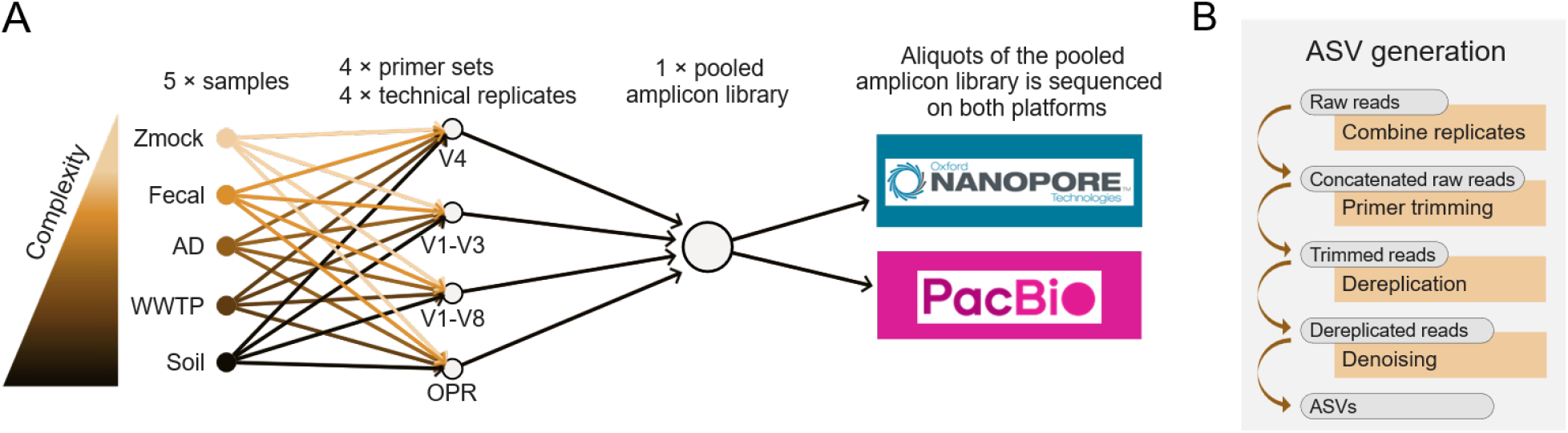
Study design and bioinformatic workflow for benchmarking ASV resolution across sequencing platforms. **A)** Samples of increasing microbial community complexity: ZymoBIOMICS mock (Zmock), ZymoBIOMICS fecal (Fecal), anaerobic digester (AD), activated sludge (WWTP), and soil communities. For each sample, amplicon libraries were created targeting regions of increasing length: the V4, V1-V3 and V1-V8 region of the 16S rRNA gene, and the full rRNA operon (OPR). Each sample-primer combination was done in four replicates. The resulting amplicon libraries were individually barcoded and pooled prior to sequencing, and the same barcoded libraries were sequenced on both Oxford Nanopore Technologies and PacBio platforms. **B)** ASVs were generated separately for each platform and primer-sample pair using a standardized bioinformatic workflow.

**Figure 2:**
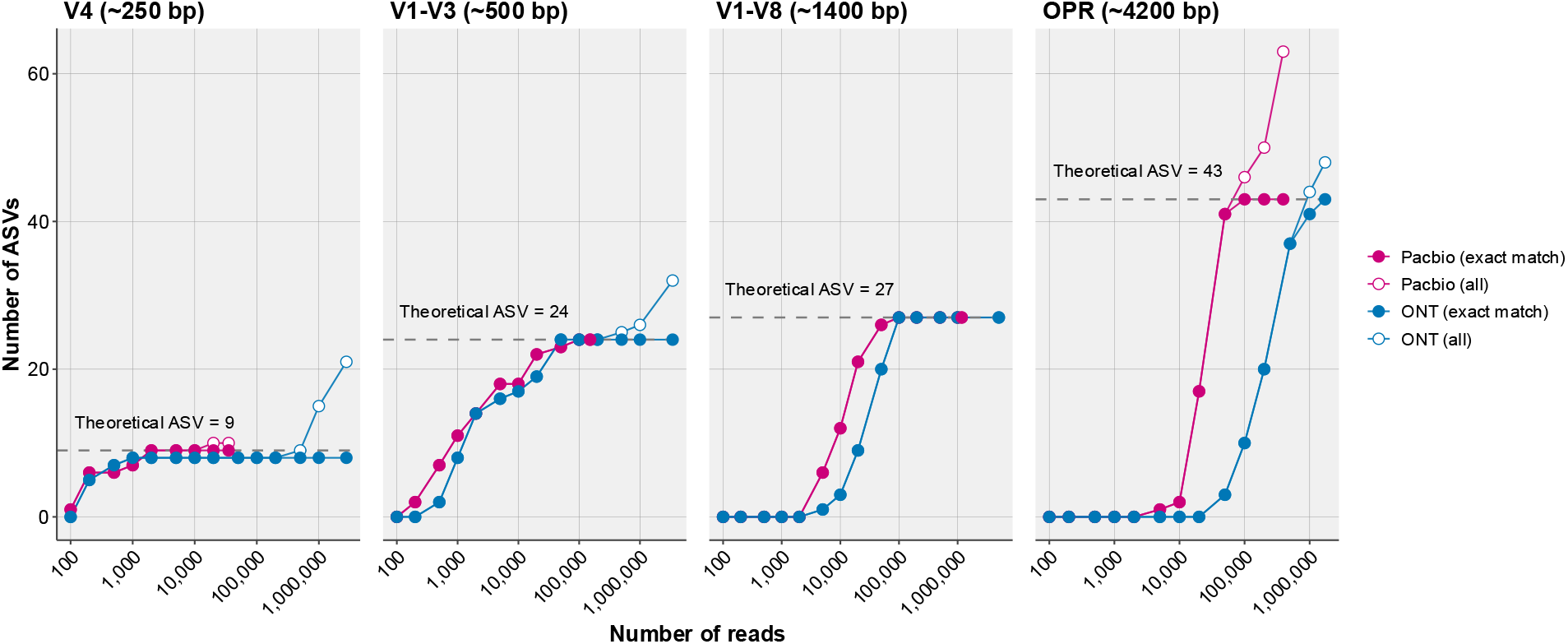
Relationship between sequencing depth and the number of called ASVs in the ZymoBIOMICS mock community for each primer set. For each sequencing platform, filled points indicate the number of ASVs with an exact match to a theoretical ASV, while open points represent the total number of ASVs. For each primer set, the dotted line marks the number of theoretical ASVs based on the curated Zmock reference sequences trimmed to match the amplified regions. Identical subsampling depths were applied across all datasets, and the final point represents the maximum number of available reads.

As the 16S gene amplicon length increased, the number of reads required to recover all sequences also increased, and this effect was substantially more pronounced for ONT. The ONT platform required 0.6×, 1.7×, 2.7×, and 5.6× more reads than PacBio for the V4, V1-V3, V1-V8 and OPR primer set, respectively (**Supplementary Fig. S1**). Together, these findings demonstrate that current ONT read quality is sufficient to generate near error-free amplicons ranging from ∼250 to >4200 bp and to resolve differences in intraspecies 16S rRNA gene diversity. When sequencing depth is adequate, ONT essentially recovers all 16S rRNA gene copy variants with perfect sequence identity. In the previous analysis, we used usearch (unoise3) for resolving ASVs. However, we obtained similar results using the DADA2 algorithm (**Supplementary Fig. S2**).

A sequencing depth of >1,000,000 reads resulted in only a few ASVs that did not match the theoretical ASVs. All non-matching ASVs were evaluated by BLAST against all Zmock reference sequences to identify the type of error, and were additionally included in the abundance-based assessment (**Supp. Table 2, Supp. Fig. S3–S5**). In V4 and V1-V3 datasets, non-matching ASVs appeared only at higher read depths, clearly indicating that the expected ASVs were generated first, as they represent the most abundant sequences. For V4 (14 sequences) and V1–V3 (8 sequences) non-matching ASVs were mostly single nucleotide polymorphisms (SNPs). These libraries were sequenced substantially deeper with ONT and in all cases these ASVs were low abundant. For OPR (21 sequences), all but one non-matching ASV could be classified as chimeras, with the remaining ASV differing by a single SNP. The chimeras were primarily formed between intragenomic copies. All but one was recovered with PacBio, and of these, four were generated by both platforms, strongly suggesting a PCR artefact rather than a sequencing error. Moreover, most non-matching ASVs were in low abundance, with only 4 of 21 being more abundant than the least abundant true ASV.

For ASV inference using usearch (unoise3), we applied the default minsize = 8, which sets the minimum abundance threshold for input sequences where lower abundances sequences are discarded prior to denoising. To evaluate the effect of relaxing this threshold, reduced minsize values were tested (**Supplementary Fig. S6**), resulting in increased ASV recovery at lower sequencing depths with correct ASVs generated first. For example, in the V1–V8 dataset, a minsize of 2 required approximately 1,730 reads to recover the first five ASVs, compared with ∼6,268 reads for minsize = 4, ∼8,442 reads for minsize = 6, and 12,647 reads for the default minsize = 8 (calculated analogously to **Supplementary Fig. S1**). At higher sequencing depths, minsize = 2 led to an increase in non-matching ASVs, likely low-abundance chimeras or sequencing artefacts, whereas minsize ≥ 4 delayed their appearance. Overall, these results indicate that moderately lowering minsize (e.g. to 4) can improve ASV recovery in complex communities, albeit with increased risk of false-positive ASVs.

To assess whether ASV inference recovers biologically meaningful abundance patterns, we examined the relative abundances of the inferred ASVs. ASVs with exact matches to the theoretical references were consistently the most abundant and closely reflected the expected 16S rRNA gene copy numbers (**Figure 3A and Supplementary Figs. S3-5A**). The abundance profiles obtained from PacBio and ONT were highly similar, and no significant differences in alpha diversity were observed between the sequencing platforms across the different primer sets (**Figure 3BC and Supplementary Figs. S3-5BC**). Differences in beta diversity were attributable to both sequencing platforms and biological replicate (n = 4); however, the overall community compositions were highly similar.

**Figure 3:**
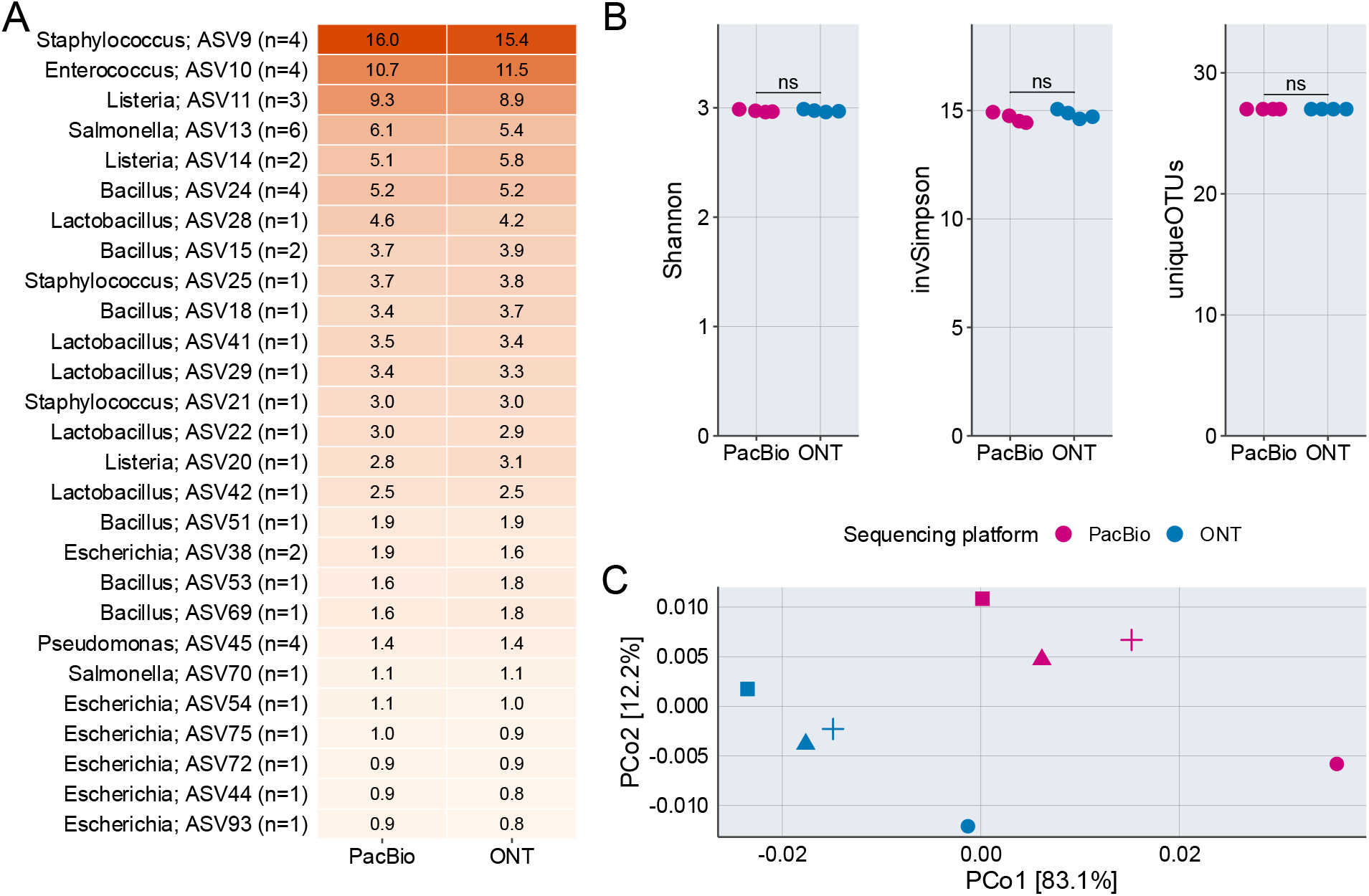
Alpha- and beta-diversity of the ZymoBIOMICS mock community using the V1-V8 primer. Data is based on all reads; however, all samples were rarefied to 228,951 reads, corresponding to the sample with the fewest reads. **A)** Relative abundance of each unique ASV, sorted by abundance for PacBio and ONT. ‘n’ indicates the copy number of the 16S rRNA gene fragment. **B)** Alpha-diversity indices; a paired t-test was performed to compare mean values between sequencing platforms (ns: not significant). **C)** Beta-diversity analysis using Bray–Curtis distances. Shapes represent biological replicates, and points are colored by sequencing platform.

Overall, our analyses confirm that ONT can recover error-free ASVs from a mock community and that the results are comparable to those obtained using PacBio.

### 16S rRNA gene ASVs can be successfully recovered from complex microbial communities

To assess the performance in real-world applications, we used four complex microbial communities representing increasing levels of diversity: a fecal sample derived from healthy human stool (fecal), a sample from an anaerobic digester (AD), a sample from an activated sludge tank at a wastewater treatment plant (WWTP), and a soil sample. ASVs were successfully recovered from all communities with both PacBio and ONT using the V4, V1-V3, and V1-V8 primer sets. However, the OPR amplicon performed poorly on ONT across all complex communities, with maximum 10 ASVs recovered (**Figure 4**).

**Figure 4:**
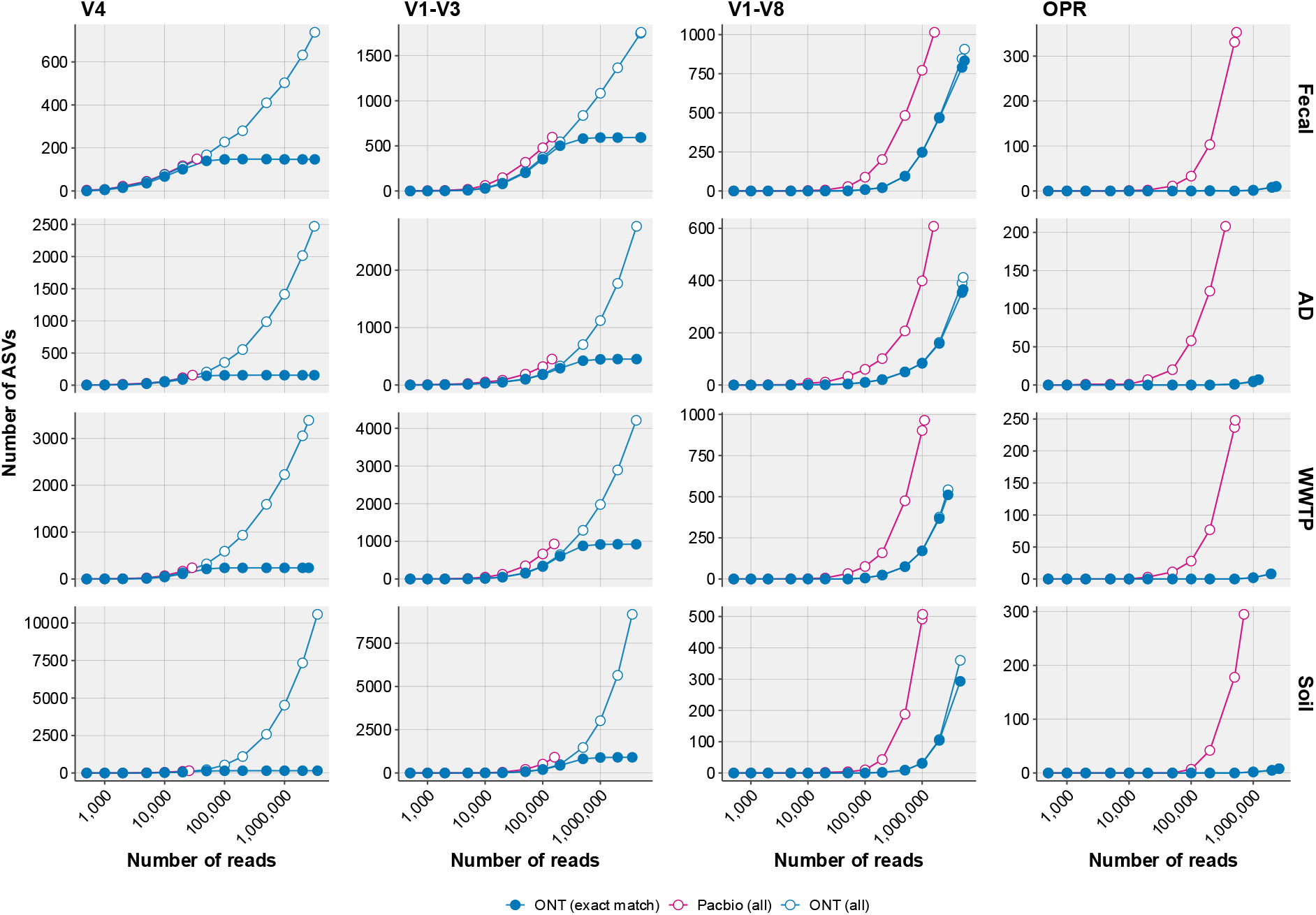
Relationship between sequencing depth and the number of called ASVs in four different complex communities for each primer set. For each sequencing platform, open points represent the total number of ASVs. Blue filled points indicate the number of ASVs from the ONT data with an exact match to an ASV recovered from the PacBio data. Identical subsampling depths were applied across all datasets, and the final point represents the maximum number of available reads. Due to low sequencing depth for PacBio V4 and V1-V3 only few subsets could be tested and a corresponding low number of ASVs were generated.

As the true composition of the complex communities is not known, we used the PacBio-derived ASVs as benchmark. Across identical subsampling depths, both platforms recovered the same ASVs and in the same order. However, we were challenged by a very different sequencing depth for especially short amplicons: Across the four replicates, PacBio yielded only ∼10,000 reads for V4 and ∼100,000 for V1-V3 due to difficulty sequencing short amplicons, whereas ONT routinely produced >1,000,000 reads. As a result, ONT generated a larger total number of ASVs simply because more reads were available. ONT recovered nearly all ASVs that were detected by PacBio (missing 0-4 ASVs for V4 and 1-8 ASVs for V1-V8 depending on sample type using all reads). PacBio retrieved the highest number of ASVs for V1-V8 and OPR primer sets. For OPR, ONT recovered only 7-10 ASVs, yet all of these were also found with PacBio. Due to the low number of generated ASV, we did not analyze the OPR data further as the number of reads required makes it unfeasible in complex communities. For V1-V8, at lower sequencing depths (∼1,000,000 reads), only 0–1 ASVs were unique to ONT, thus the first >300 ASV - likely the most abundant - were shared across platforms. At higher sequencing depths (i.e. using all reads), differences between platforms became more pronounced, with 31-74 ASVs unique to ONT. This pattern is expected, as increasing sequencing depth progressively samples the rare biosphere (less abundant ASVs), where ASV detection is increasingly influenced by stochastic sampling effects rather than consistent biological signal [22].

Together, these results indicate that ONT-unique ASVs in long-amplicon datasets likely represent genuine community diversity rather than sequencing artifacts, and that reliable ASV inference can be achieved using ONT reads for long amplicons such as V1-V8, though it requires higher read depth to match PacBio’s recovery. To obtain 100 ASVs, ONT needed approximately 2-3× more reads for V4 and V1-V3, 4.1-5.6× more reads for V1-V8, and 25-42× more reads for OPR (**Supplementary Fig S7**). This was consistent with observations from the Zmock, however with larger fold difference for the complex communities. We hypothesized that this effect was due to higher ONT basecalling accuracy for well-characterized reference strains, which would likely have been used to train the basecalling model. We therefore compared the average read quality for each sample after primer filtration. In support of our hypothesis, we found that the average Phred-score was approximately one unit lower in the complex communities than in the corresponding Zmock dataset (**Supplementary Fig. S8**).

Next, we examined the obtained microbial profiles for the different complex environments and primer sets (**Figure 5 and Supplementary Figs. S9-11**). For V1-V8, where we had the most data, differences in alpha diversity indexes between sequencing platforms were either insignificant or minor. Examining the most abundant ASVs per community, relative abundances were nearly identical, with fecal samples showing the largest discrepancy, consistent with the variation observed in alpha diversity indexes. Beta-diversity patterns were assessed for each primer set and sample type. Although both sequencing platform and biological replicate contributed to the observed variation, samples exhibited highly similar community compositions (**Supplementary Fig. S12**). This clearly illustrates that we were able to obtain highly similar communities in terms of both alpha- and beta-diversity with ONT and PacBio, using an ASV-based framework. This finding contrasts with previous benchmarking studies, where differences in alpha and beta diversity were reported when comparing ONT alignment or OTU-based approaches with PacBio or Illumina ASV-based workflows [23–25].

**Figure 5:**
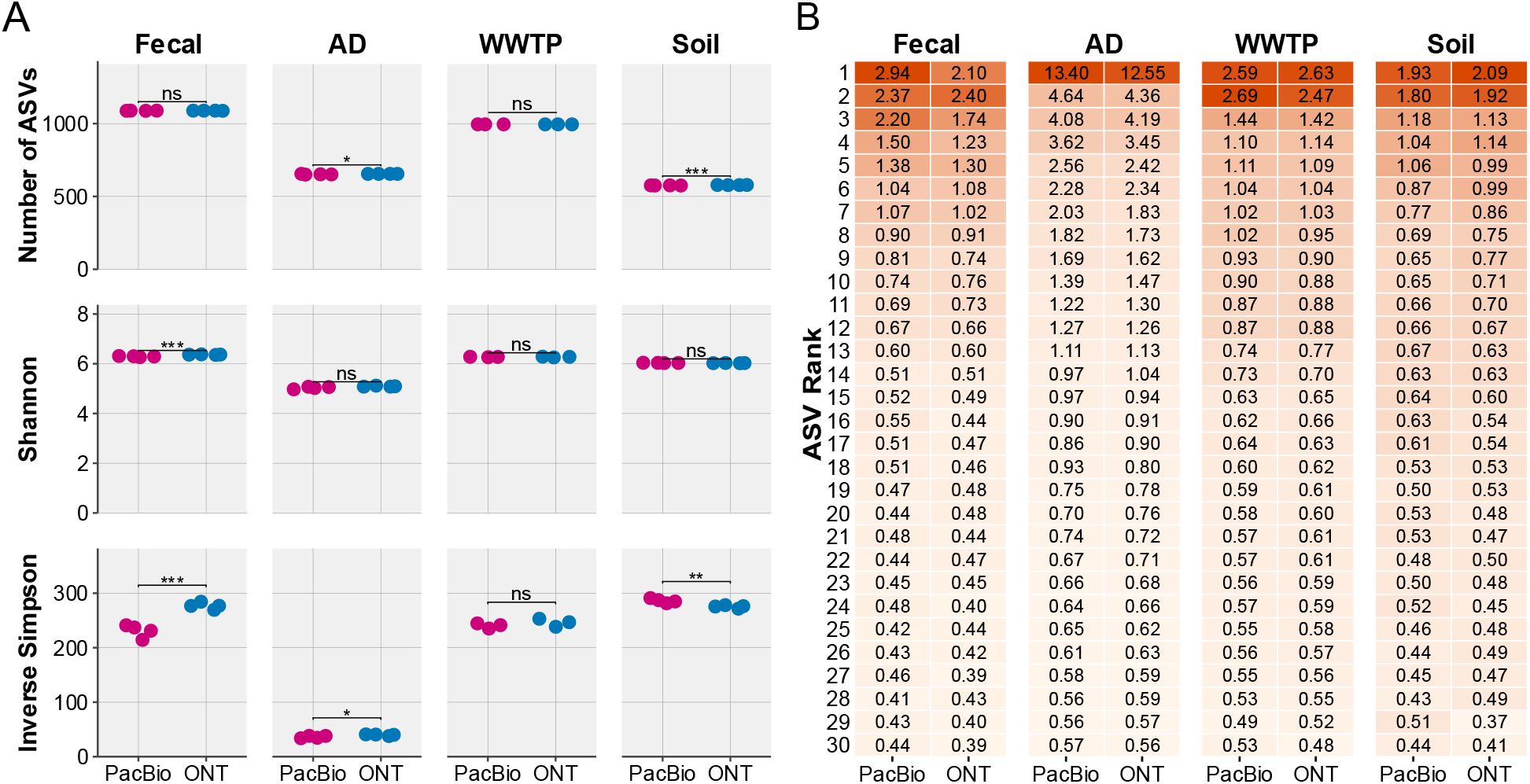
Alpha-diversity and top 30 most abundant ASVs in the complex communities (V1-V8 data). **A)** Alpha-diversity indices; a t-test was performed to compare mean values between sequencing platforms (ns: not significant; *: p<0.05; **: p<0.01; ***: p<0.001). **B)** Relative abundance of the top 30 most abundant ASVs in each sample type with the relative abundance shown in percent. All analyses were performed on amplicon data rarefied to 88,500 reads, corresponding to the sample with the fewest reads.

Finally, we evaluated the effects of primer choice and sequencing platform on community-level patterns at the family level (**Figure 6**). Analyses were conducted at this level, as the V4 provides limited taxonomic resolution. The bias introduced by primer selection was consistently larger than the effect of sequencing platform, and only for the fecal community a significant difference was observed between sequencing platform (*R*^2^=0.22, *P*<0.003), while the choice of primer set was significant for all, explaining 55% (fecal) to 88% (soil) of the community variation (*P*<0.001). To assess whether the observed trends persisted at higher sequencing depth and taxonomic resolution, the same analyses were repeated using only the V1-V3 and V1-V8 datasets at genus level. This yielded the same result, however with primer set explaining 86-97% and sequencing platform insignificant for all sample types (**Supplementary Fig. S13**).

**Figure 6:**
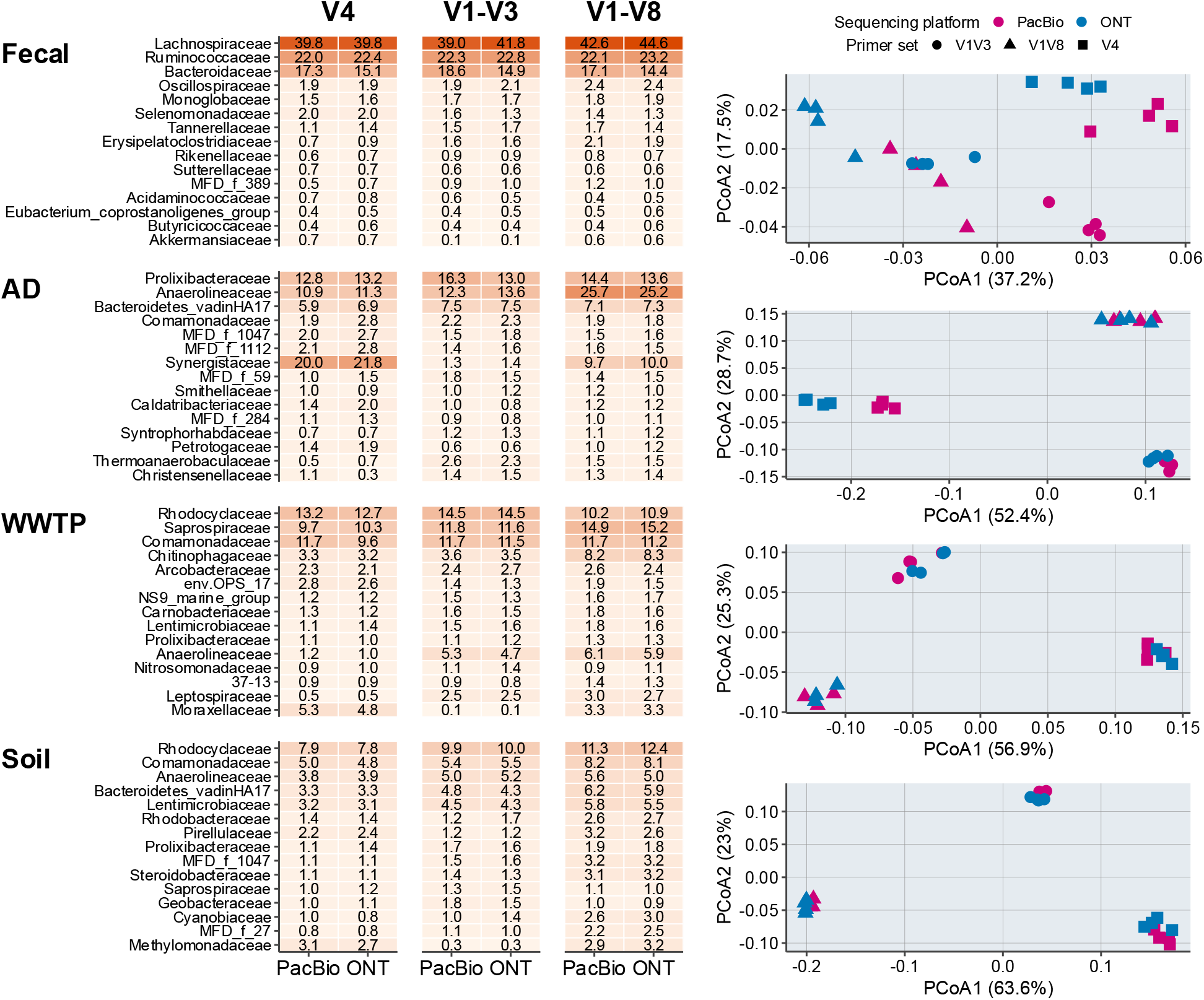
Taxonomic diversity across primer sets and sample types. All analyses were performed on rarefied datasets, with read counts normalized to the sample with the fewest reads (5,126 reads). **A)** Top 15 most abundant families across each sample content. **B)** Beta diversity of samples at the family level for each sample type based on Bray-Curtis distance matrices, including only ASVs classified to the family level and with a relative abundance ≥ 0.1%. PERMANOVA results were assessed for the effects of sequencing platform (SP) and primer set (PS). Fecal: SP: *R*^2^=0.22, *P*<0.003, PS: *R*^2^=0.55, *P*<0.001; AD: SP: *R*^2^=0.03, *P*<0.516, PS: *R*^2^=0.86, *P*<0.001; WWTP: SP: *R*^2^=0.02, *P*<0.844, PS: *R*^2^=0.82, *P*<0.001; Soil: SP: *R*^2^=0, *P*<0.983, PS: *R*^2^=0.88, *P*<0.001.

Our results demonstrate that the ONT platform enables robust ASV inference from complex microbial communities, consistently recovering all abundant ASVs also detected using PacBio. Alpha-diversity estimates were highly concordant between platforms, contrasting with most previous benchmarking studies that report substantial platform-dependent biases. Across all analyses, community composition profiles were highly similar between sequencing technologies, with primer choice accounting for substantially more variation than sequencing platform, demonstrating that platform-specific effects are minimal.

## Conclusion

We demonstrate that *de novo* ASV-resolved profiling can now be achieved using the ONT platform for amplicon sequencing. Across mock communities, ONT data resolved intragenomic 16S rRNA gene variants using V4, V1-V3, V1-V8, and full-length operon (OPR) amplicons spanning ∼250-4,200 bp. For complex communities, we recovered the same ASVs with ONT as with PacBio; however, greater sequencing depth was required for ONT: approximately 2-3× more reads for V4 and V1-V3, 4.1-5.6× more reads for V1-V8, and 25-42× for full-length operon primers. Consequently, using OPR primers we generated ≤ 10 ASVs with ONT, limiting validation with the PacBio-derived ASVs and showing that sequencing long amplicons (∼4,200 bp) from complex communities is currently not cost-effective with ONT. Adjusting the default abundance threshold (minsize) increased ASV recovery from smaller subsets of reads, thus improving ASV resolution for longer amplicons (e.g., V1-V8) and potentially enhancing the feasibility of OPR-based profiling, albeit with an increased risk of false positives. When sufficient sequencing depth was achieved, ONT accurately recovered the same ASVs with perfect sequence identity to ASVs derived from PacBio data. Community profiles generated from ONT and PacBio data were highly concordant in both alpha- and beta-diversity metrics. This concordance held when analyses were performed at the ASV level within individual primer sets and communities, as well as across primer sets, with primer choice explaining most community variation and sequencing platform contributing only a minor effect. Together, these results demonstrate that ONT amplicon sequencing has matured beyond OTU-based or reference-dependent workflows, enabling true ASV-based profiling even in environments lacking comprehensive reference databases.

## Materials and Methods

### Sample collection and DNA extraction

We collected five samples for analysis: A soil sample from the surroundings of a public pond in Aalborg SØ (collected 2025-07-21), an anaerobic digester (AD) sample from Randers wastewater treatment plant (collected 2016-03-07), an activated sludge sample from Aalborg West wastewater treatment plant (WWTP) (collected 2025-05-27) and two synthetic microbial communities: ZymoBIOMICS Microbial Community DNA Standard (cat. no. D6305, Zymo Research, USA) and ZymoBIOMICS Gut Microbial Standard (cat. no. D6323, Zymo Research, USA). DNA from the soil sample was extracted using the QIAGEN DNeasy PowerSoil Pro Kit. From WWTP and AD, DNA was extracted following the corresponding standard MiDAS protocols (https://www.midasfieldguide.org/guide/protocols). DNA purity, concentration and size was assessed with NanoDrop OneC (Thermo Scientific), Invitrogen’s Qubit 1X dsDNA HS kit on a Qubit 4.0 fluorometer, and Agilent’s Genomic DNA ScreenTape Analysis.

### 16S rRNA gene and operon amplicon sequencing

Barcoded amplicons were generated in four replicates per sample-primer combination using a two-step PCR approach, in which target regions were first amplified with locus-specific primers carrying ONT universal tag overhangs and subsequently barcoded in a second PCR using the Nanopore EXP-PBC096 barcoding kit (**Table 1**).

**Table 1:**
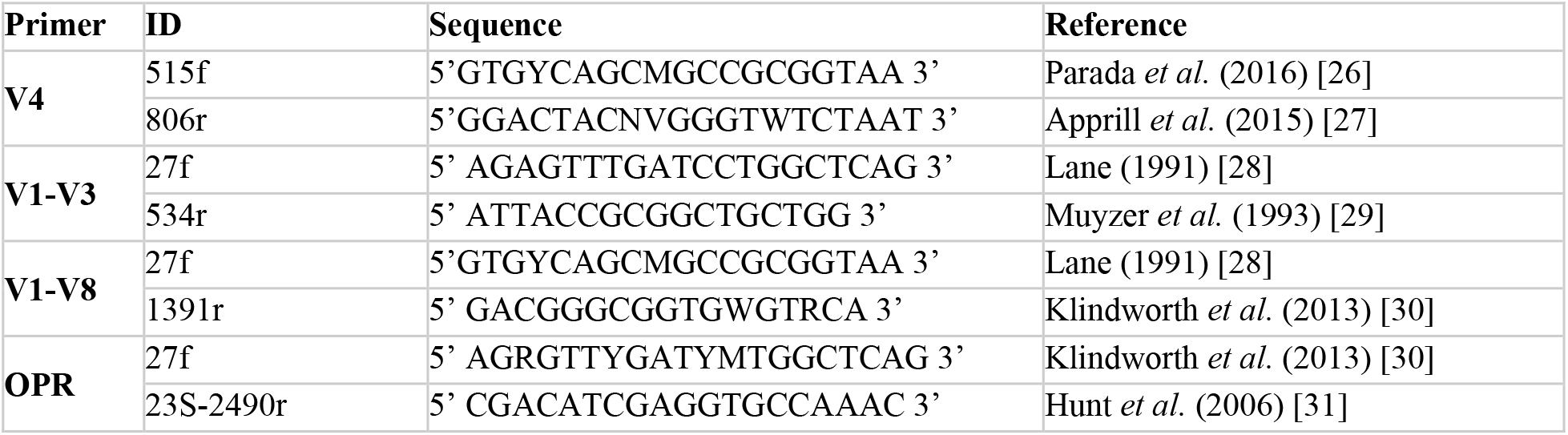
Overview of primers used in this study. All primers used for PCR contained a 5’ ONT universal tag overhang for subsequent barcoding: 5’-TTTCTGTTGGTGCTGATATTGC-3’ for forward primers and 5’-ACTT-GCCTGTCGCTCTATCTTC-3’ for reverse primers.

For V1-V3, V4 and V1-V8, the first PCR reaction contained 4 µl of 5ng/µl DNA, 200 nM of each primer, 25 µl of PCRBIO 2X UltraMix and nuclease free water in a final volume of 50 µl. Reactions were split into two reactions of 25 µl and combined after amplification was finished. The thermocycler settings for amplification was as follows: Initial denaturation at 95°C for 2 minutes followed by 20 (V1-V3) or 25 (V4 and V1-V8) cycles of denaturation at 95°C for 15 seconds, annealing at 55°C for 15 seconds, and extension at 72°C for 50 (V1-V3 and V4) or 120 (V1-V8) seconds, followed by a final extension at 72°C for 5 minutes.

The PCR reactions for the rRNA operon (OPR) were carried out with 4 µl of 5 ng/µl DNA, 25 µl of NEBNext Ultra II Q5 MasterMix, 500 nM of each primer, and nuclease-free water in a final volume of 50 µl. The thermocycler settings for amplification were as follows: Initial denaturation at 98°C for 30 seconds, followed by 25 cycles of denaturation at 98°C for 10 seconds, annealing at 55°C for 15 seconds, and extension at 72°C for 2 minutes, followed by a final extension at 72°C for 2 minutes.

The resulting amplicons were purified by adding AmPure XP bead solution in a ratio of 0.8x bead/sample for the V4, V1-V3 and V1-V8 regions and 0.7x bead/sample for the OPR amplicons, incubated at room temperature for 5 minutes and washed twice in 80% ethanol, and finally eluted in 20 µl of nuclease-free water. Concentration and quality of the obtained PCR products was assessed using the Qubit dsDNA HS kit in a Tecan Infinite 200 Pro Microplate reader, and an Agilent D1000 (V1-V3 and V4) or D5000 (V1-V8 and OPR) ScreenTape.

Barcoding of amplicons was carried out in 50 µl reactions containing 200 fmol of amplicons, 1 µl of PCR barcode, and 25 µl of LongAmp Taq 2x master mix. The thermocycler settings for amplification was as follows: Initial denaturation at 95°C for 3 minutes, followed by 12 cycles of denaturation at 95°C for 15 seconds, annealing at 62°C for 15 seconds, and extension at 65°C for 60 s (V1-V3, V4, and V1-V8) or 4 m (OPR), followed by a final extension at 65°C for 5 (V1-V3, V4, and V1-V8) or 8 (OPR) minutes. PCR products were purified and concentrations measured as mentioned above with AmPure XP bead solution in a ratio of 0.8x bead/sample. Libraries were pooled in equivalent molar concentrations, and concentration and integrity of the resulting pool was verified with the Qubit dsDNA HS kit on a Qubit 4.0 fluorometer, and Agilent D5000 DNA ScreenTape.

For nanopore sequencing, 800 ng of the pooled amplicon library prepared for sequencing using the ligation sequencing kits SQK-LSK114 (Oxford Nanopore Technologies) and sequenced on a PromethION flowcells (FLO-PRO114M) on a PromethION P2Solo. Basecalling was carried out using dorado (v.1.0.0) using the super accuracy model (v.5.2.0). For PacBio sequencing, 300 ng of the pooled amplicon library was converted to SMRTbell libraries using the SMRTbell prep kit 3.0 and sequenced on a Revio SMRT Cell 25M using a 24-hour movie and aiming for an on-plate loading concentration of 300 pM (based on 3000 bp as average insert size) on a PacBio Revio instrument.

### Bioinformatic processing of amplicon data

The following standard workflow primary based on USEARCH v11.0.667 [6] was used for processing the amplicon data unless otherwise stated: Raw reads from both PacBio and ONT were demultiplexed using dorado v1.1.1 (--kit-name EXP-PBC096), relabeled based on sample ID using usearch -fastx_relabel -prefix, and reads from samples, which should be analyzed together were concatenated into combined fastq files. Primer trimming was carried out with Cutadapt v.5.1 [32] using the following parameters -g {Fw…RCRev} --revcomp --discard-untrimmed, where Fw and RCRev are the forward primer and reverse complement reverse primer without overhangs, respectively, found in **Table 1**. Empty sequences generated during primer trimming were removed using a custom script. Reads were dereplicated using usearch -fastx_uniques -minuniquesize 2 - sizeout. ASVs were resolved using usearch -unoise3 -minsize 8. ASVs were further filtered by expected amplicon length, defined as ±25% of the median length of the corresponding ZymoBIOMICS reference sequences (**Supplementary Fig. S14**). ASV-tables were created mapping the primer-trimmed reads against the ASVs using usearch -otutab -zotus.

For each primer set and sample content, ASVs generated from PacBio and ONT were merged prior to mapping, producing count tables for downstream alpha- and beta-diversity analyses. Taxonomy was assigned to the ASVs using the SINTAX classifier [33] with usearch -sintax -strand plus - sintax_cutoff 0.8 using the MicroFlora Global 16S rRNA gene reference database v.1.3 [34].

To evaluate the impact of ASV-calling algorithm, we also resolved ASVs DADA2 [35]. This was done in qiime2 v.2025.10 [36] using the following command: dada2 denoise-single and standard settings.

To test how the read depth affected our ability to resolve ASVs, merged reads for each combination of sample, primer set, and sequencing platform was subsampled to various depths after primer-trimming using usearch -fastx_subsample. For analysis of the ZymoBIOMICS mock community, curated reference sequences were obtained from Karst *et al*. (2021) [16] and Lin *et al*. (2024) [18] and the sequences were trimmed with Cutadapt and demultiplexed to generate primer-specific reference amplicons that were mapped to the generated ASVs using usearch -search_exact (**Supplementary file 1**).

All downstream statistical analyses, diversity analyses, and data visualization were performed in R v.4.4.0, [37] using RStudio Server (v.2024.04.2+764) [38]. Analyses were conducted primarily with the tidyverse packages [39], together with additional packages including ampvis2 and vegan [40, 41].

## Data and Code Availability

The raw sequencing data generated in this study have been deposited in the Sequence Read Archive (SRA) at NCBI under accession number PRJNA1389832. Scripts for ASV generation, postprocessing, and figures are available at https://github.com/MarieRiisgaard/ont-asv-microbial-profiling.

## Acknowledgement

The project has been funded by the Novo Nordisk Foundation (REThiNk, grant NNF22OC0071498, P.H.N. and M.K.D.D).

