## Supplementary tables and figures for "Nanopore sequencing reaches amplicon sequence variant (ASV) resolution"

**Supplementary Table S1: Composition of the ZymoBIOMICS Microbial Community DNA Standard (Zmock) (Cat. No. D6305, Zymo Research, USA)**. Species composition and genomic DNA abundance were adapted from datasheet provided by ZymoBIOMICS (D6305), and 16S rRNA gene copy numbers were obtained from Lin et al. (2024)*

| Species | Avg. GC (%) | Gram stain | gDNA abundance (%) | 16S copy number |
|---|---|---|---|---|
| *Pseudomonas aeruginosa* | 66.2 | − | 12 | 4 |
| *Escherichia coli* | 56.8 | − | 12 | 7 |
| *Salmonella enterica* | 52.2 | − | 12 | 7 |
| *Lactobacillus fermentum* | 52.8 | + | 12 | 5 |
| *Enterococcus faecalis* | 37.5 | + | 12 | 4 |
| *Staphylococcus aureus* | 32.7 | + | 12 | 6 |
| *Listeria monocytogenes* | 38.0 | + | 12 | 5 |
| *Bacillus subtilis* | 43.8 | + | 12 | 9 |
| *Saccharomyces cerevisiae* | 38.4 | Yeast | 2 | NA |
| *Cryptococcus neoformans* | 48.2 | Yeast | 2 | NA |

*Lin, X., Waring, K., Ghezzi, H., Tropini, C., Tyson, J., Ziels, R.M., 2024. High accuracy meets high throughput for near full-length 16S ribosomal RNA amplicon sequencing on the Nanopore platform. PNAS Nexus 3, pgae411. https://doi.org/10.1093/pnasnexus/pgae411

**Supplementary Table S2: Evaluation of non-matching ASVs compared to the Zmock reference sequences.** The table includes ASVs generated using the unoice3 algorithm. Each ASV is reported with its ASV ID and the primer used (OPR: full rRNA operon primers). Platform indicates the sequencing platform, with PB denoting PacBio and PB:ONT indicating that the ASV was identified using both sequencing platforms. ASVs were evaluated by using BLAST of all ASVs against all reference sequences to identify the type of error, noted in the comment column. SNP: single-nucleotide polymorphism.

| ASV ID | Primer | Platform | Reference match | Error type | Comment |
|---|---|---|---|---|---|
| **ASV1029** | OPR | PB | Non-exact | Chimera | Between different genera |
| **ASV1306** | OPR | PB | Non-exact | Chimera | Between different genera |
| **ASV1363** | OPR | PB | Non-exact | Chimera | Between intragenomic copies |
| **ASV1372** | OPR | PB | Non-exact | Chimera | Between intragenomic copies |
| **ASV1525** | OPR | PB | Non-exact | Chimera | Between intragenomic copies |
| **ASV1535** | OPR | PB | Non-exact | Chimera | Between intragenomic copies |
| **ASV1539** | OPR | PB | Non-exact | SNP | TT->AA |
| **ASV1578** | OPR | PB | Non-exact | Chimera | Between different genera |
| **ASV1580** | OPR | PB | Non-exact | Chimera | Between different genera |
| **ASV1581** | OPR | PB | Non-exact | Chimera | Between intragenomic copies |
| **ASV1582** | OPR | PB | Non-exact | Chimera | Between intragenomic copies |
| **ASV1586** | OPR | PB | Non-exact | Chimera | Between intragenomic copies |
| **ASV209** | OPR | PB;ONT | Non-exact | Chimera | Between intragenomic copies |
| **ASV463** | OPR | ONT | Non-exact | Chimera | Between intragenomic copies |
| **ASV544** | OPR | PB;ONT | Non-exact | Chimera | Between intragenomic copies |
| **ASV558** | OPR | PB | Non-exact | Chimera | Between intragenomic copies |
| **ASV571** | OPR | PB;ONT | Non-exact | Chimera | Between different genera |
| **ASV657** | OPR | PB;ONT | Non-exact | Chimera | Between intragenomic copies |
| **ASV773** | OPR | PB | Non-exact | Chimera | Between intragenomic copies |
| **ASV962** | OPR | PB | Non-exact | Chimera | Between intragenomic copies |
| **ASV965** | OPR | PB | Non-exact | Chimera | Between intragenomic copies |

| | | | | | |
|---|---|---|---|---|---|
| **ASV10128** | V1-V3 | ONT | Non-exact | SNP | 3G->2G+CT->TC |
| **ASV10228** | V1-V3 | ONT | Non-exact | SNP | 3G->2G |
| **ASV13820** | V1-V3 | ONT | Non-exact | SNP | 3G->2G+CT->TC (both same as for V1V3; ASV10128) |
| **ASV13927** | V1-V3 | ONT | Non-exact | SNP | Gap->A+T->G (same as for V1V3; ASV7035) |
| **ASV15068** | V1-V3 | ONT | Non-exact | SNP | 4G->3G |
| **ASV15071** | V1-V3 | ONT | Non-exact | SNP | 3G->2G (same as for V1V3;ASV10128) |
| **ASV7035** | V1-V3 | ONT | Non-exact | SNP | Gap->A+T->G |
| **ASV7642** | V1-V3 | ONT | Non-exact | SNP | 3G->2G (same as for V1V3; ASV10128) |
| **ASV10441** | V4 | ONT | Non-exact | SNP | 6G-5G |
| **ASV10444** | V4 | ONT | Non-exact | SNP | 6G->5G |
| **ASV10445** | V4 | ONT | Non-exact | SNP | 3G->2G |
| **ASV10446** | V4 | ONT | Non-exact | Chimera | Between different genera |
| **ASV10718** | V4 | ONT | Non-exact | Chimera | Between intragenomic copies |
| **ASV10721** | V4 | ONT | Non-exact | Chimera | Between different genera |
| **ASV15073** | V4 | ONT | Non-exact | SNP | A->T+2T->T |
| **ASV3562** | V4 | ONT | Non-exact | SNP | 4G->3G |
| **ASV6530** | V4 | ONT | Non-exact | Chimera | Between different genera |
| **ASV6926** | V4 | ONT | Non-exact | Conta-minant | Perfect hit to *Paenibacillus cookii* sequence in NCBI |
| **ASV7004** | V4 | PB | Non-exact | Chimera | Between different genera |
| **ASV7074** | V4 | ONT | Non-exact | SNP | 2T->T |
| **ASV9469** | V4 | ONT | Non-exact | SNP | 5C->4C |
| **ASV9486** | V4 | ONT | Non-exact | Chimera | Between different genera |

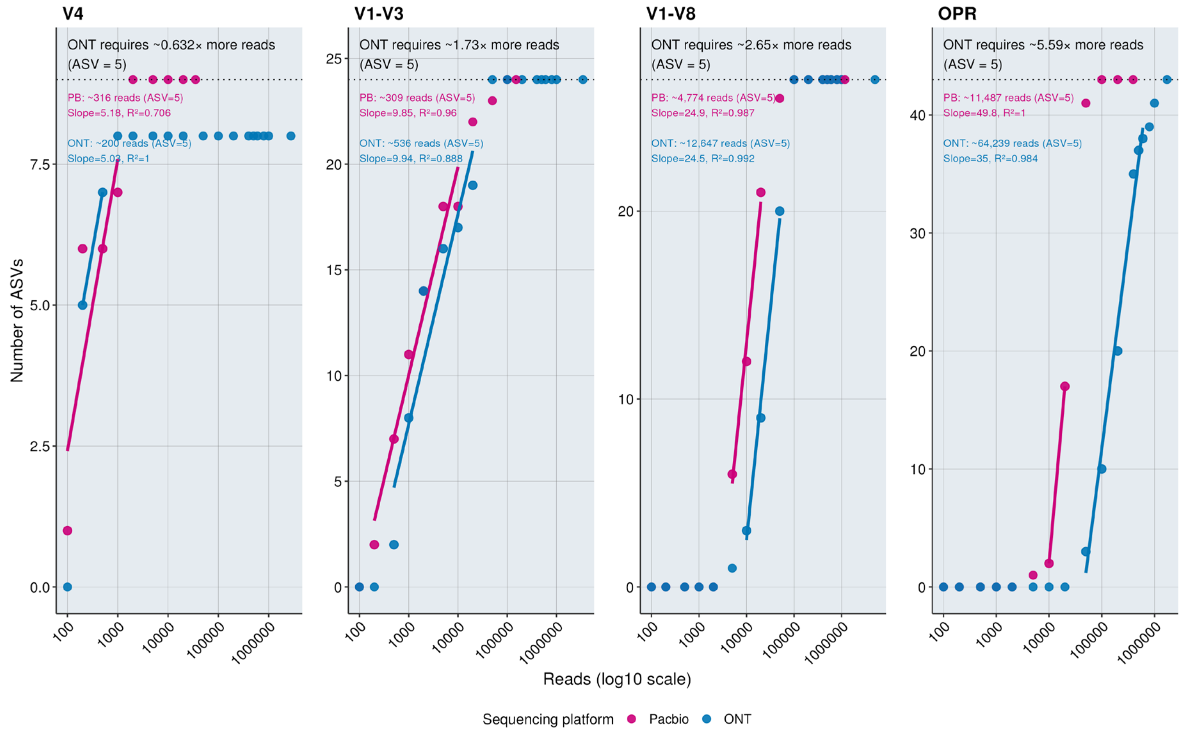


**Supplementary Fig. S1: ASV accumulation curves and linear-phase efficiency for PacBio and ONT across primer sets.** For each primer set (V4, V1-V3, V1-V8, and OPR), the number of recovered ASVs is shown as a function of sequencing depth for PacBio and ONT. Linear-phase points were identified using primer-specific criteria, and linear models of ASV count versus log10-transformed read depth were fitted to this region only (solid lines). The horizontal dotted line marks the expected number of true ASVs in the mock community. For each platform, the panel reports: (i) the estimated read depth required to recover 5 ASV, (ii) the slope and $R^2$ of the linear-phase model, and (iii) the fold-difference in reads required between ONT and PacBio**.**

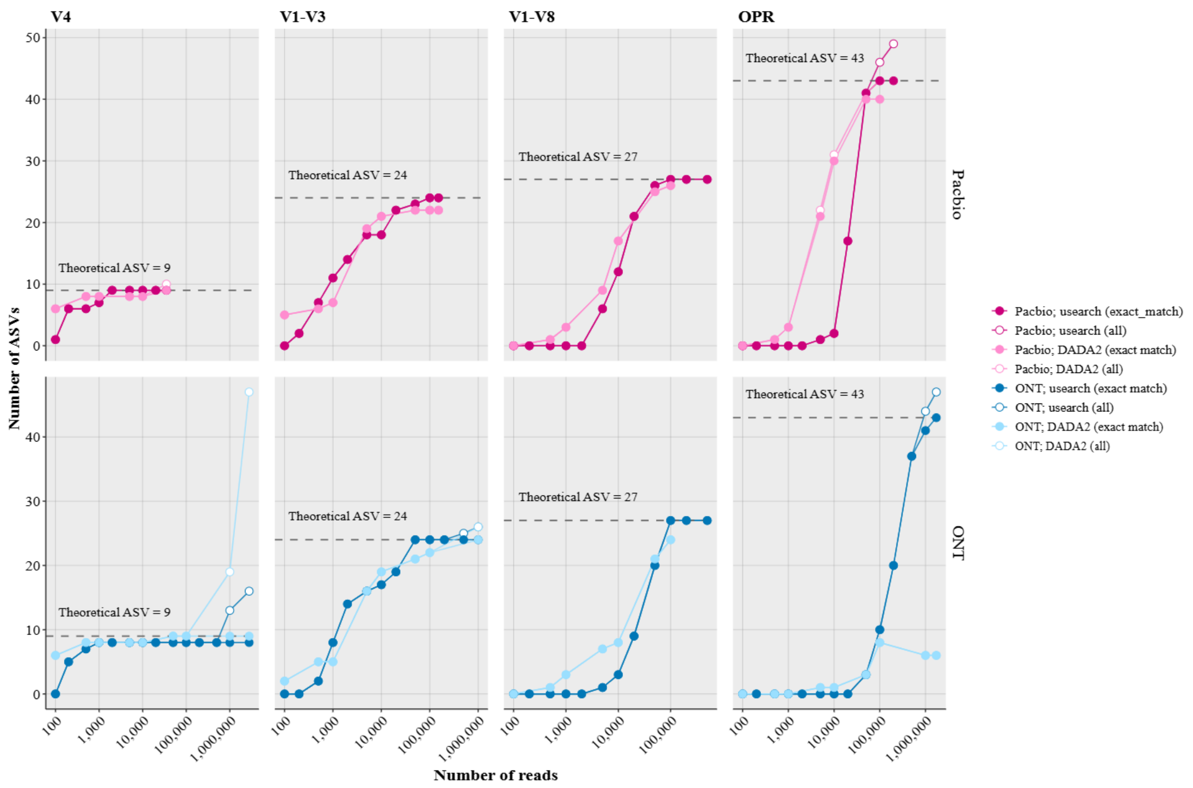


**Supplementary Fig. S2: Relationship between sequencing depth and the number of called ASVs in the ZymoBIOMICS mock community for each primer set, using either the DADA2 or UNOISE algorithm for denoising**. For each sequencing platform, filled points indicate the number of ASVs with an exact match to a theoretical ASV, while open points represent the total number of ASVs after length and chimera filtering. For each primer set, the dotted line indicates the number of theoretical ASVs based on curated Zmock reference sequences. Identical subsampling depths were applied across all datasets, and the final point represents the maximum number of available reads. All subsamples were processed with both DADA2 and UNOISE; however, some DADA2 data points are missing because the algorithm did not complete successfully for those subsamples.

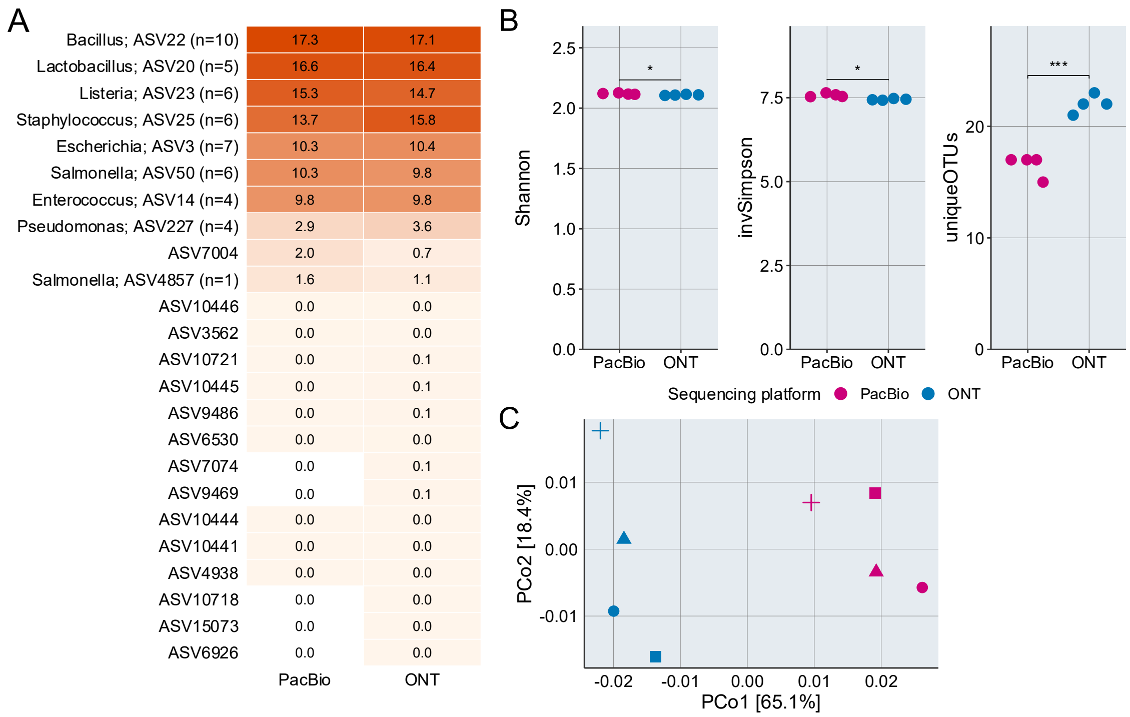


**Supplementary Fig. 3: Alpha- and beta-diversity of the ZymoBIOMICS mock community using the V4 primer.** Data is based on all reads; however, all samples were rarefied to 7,173 reads, corresponding to the sample with the fewest reads. **A)** Relative abundance of each unique ASV, sorted by abundance, for PacBio and ONT. 'n' indicates the copy number of the 16S rRNA gene fragment. A dot indicates ASVs that were filtered due to length constraints or identified as chimeric sequences. **B)** Alpha-diversity indices; a paired t-test was performed to compare mean values between sequencing platforms (ns: not significant; *: $P<0.05$; **: $P<0.01$; ***: $P<0.001$). **C)** Beta-diversity analysis using Bray-Curtis distances. Shapes represent biological replicates.

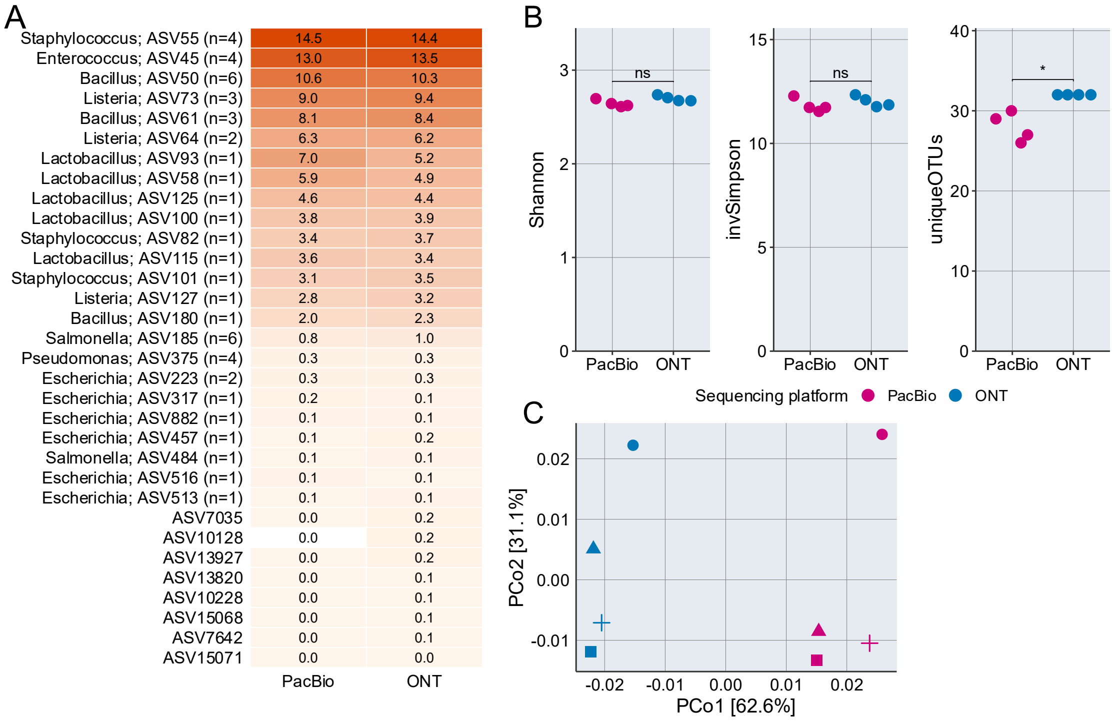


**Supp. Fig. 4: Alpha- and beta-diversity of the ZymoBIOMICS mock community using the V1-V3 primer.** Data are based on all reads; however, all samples were rarefied to 24,668 reads, corresponding to the sample with the fewest reads. **A)** Relative abundance of each unique ASV, sorted by abundance, for PacBio and ONT. 'n' indicates the copy number of the 16S rRNA gene fragment. A dot indicates ASVs that were filtered due to length constraints or identified as chimeric sequences. **B)** Alpha-diversity indices; a paired t-test was performed to compare mean values between sequencing platforms (ns: not significant; *: *P*<0.05; **: *P*<0.01; ***: *P*<0.001). **C)** Beta-diversity analysis using Bray-Curtis distances. Shapes represent biological replicates.

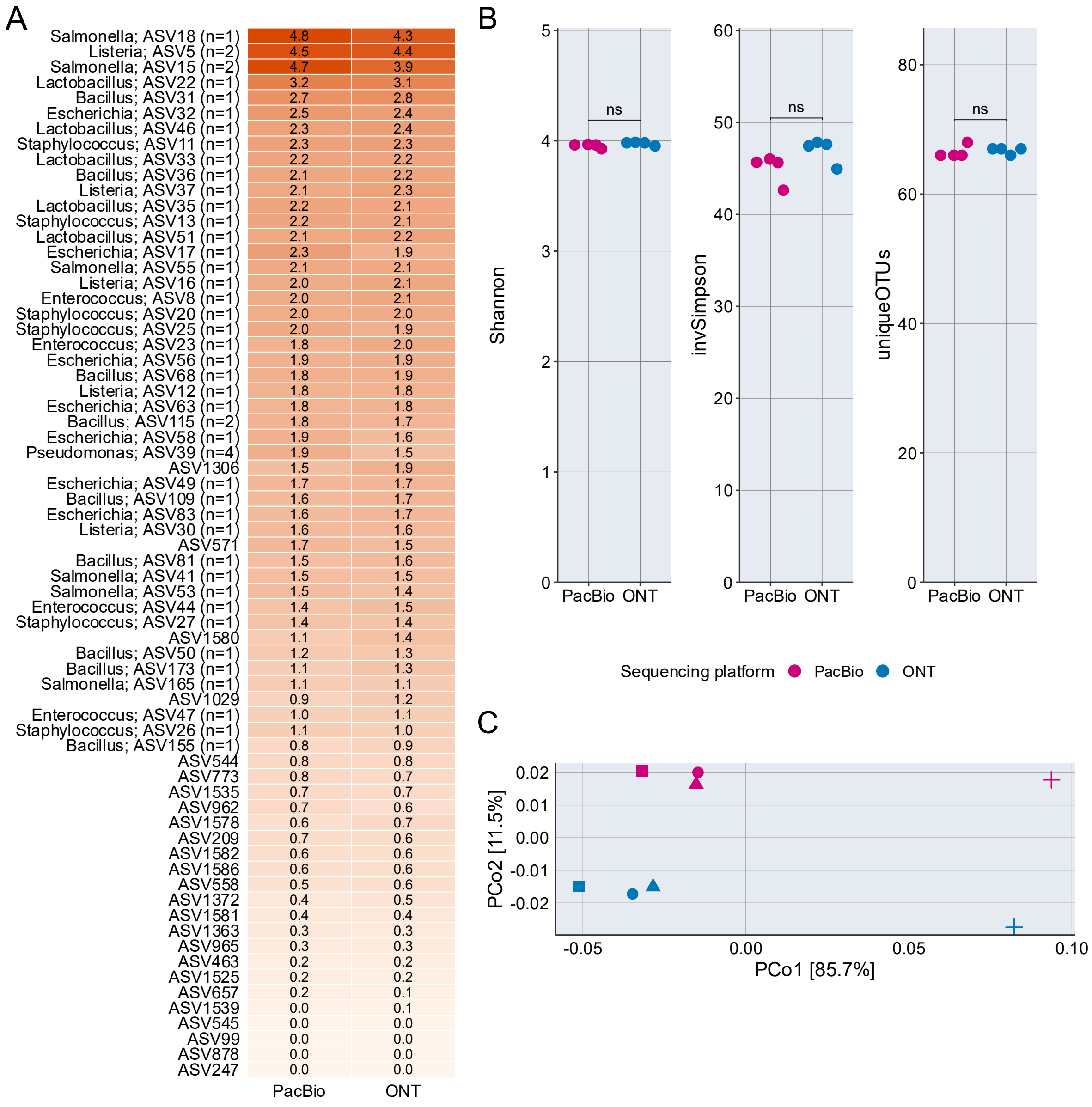


**Supp. Fig. 5: Alpha- and beta-diversity of the ZymoBIOMICS mock community using the full rRNA operon primers (OPR).** Data is based on all reads; however, all samples were rarefied to 54,149 reads, corresponding to the sample with the fewest reads. **A)** Relative abundance of each unique ASV, sorted by abundance, for PacBio and ONT. 'n' indicates the copy number of the 16S rRNA gene fragment. A dot indicates ASVs that were filtered due to length constraints or identified as chimeric sequences. **B)** Alpha-diversity indices; a paired t-test was performed to compare mean values between sequencing platforms (ns: not significant; *: $P<0.05$; **: $P<0.01$; ***: $P<0.001$). **C)** Beta-diversity analysis using Bray-Curtis distances. Shapes represent biological replicates.

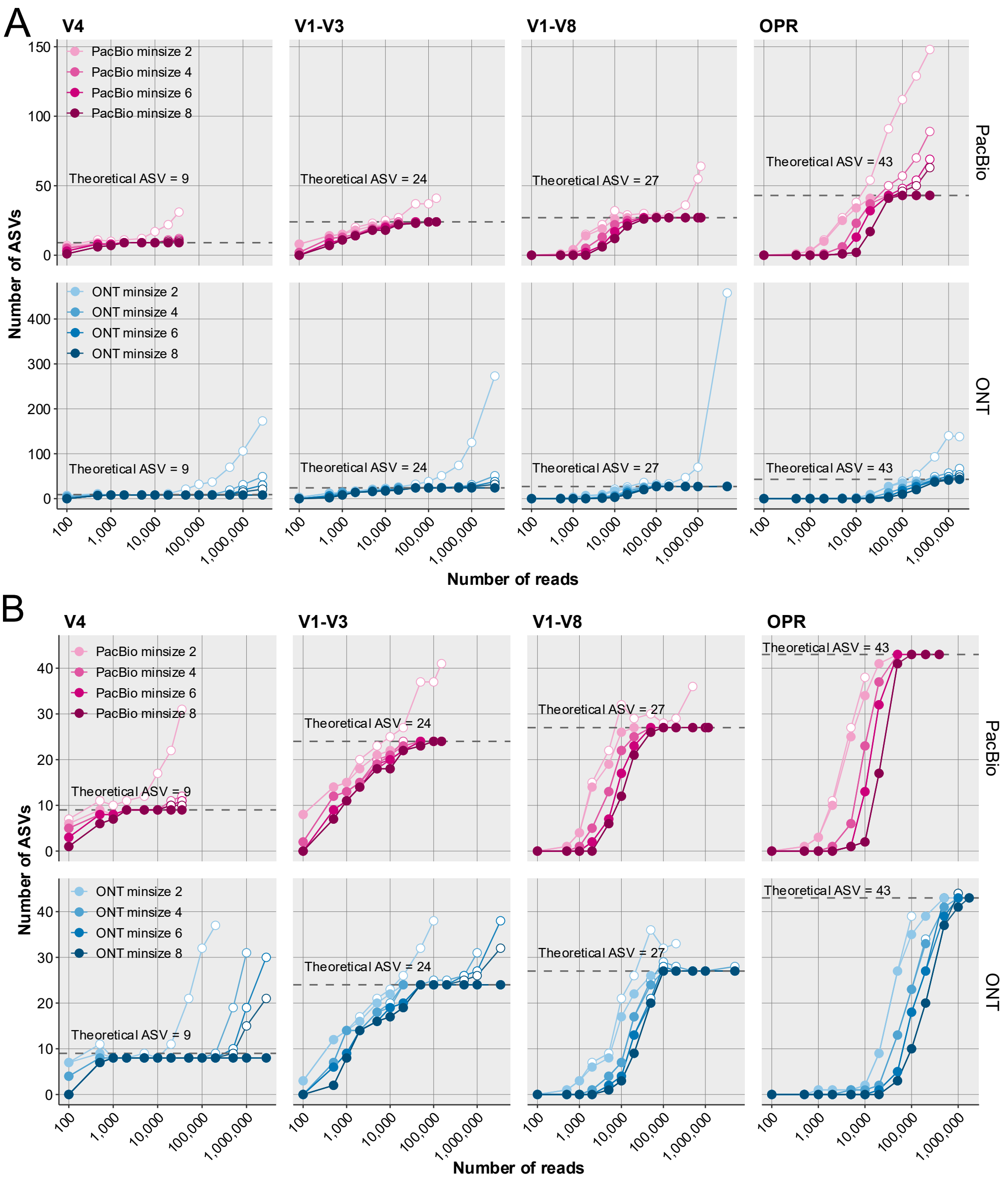


**Supp. Fig. 6: Relationship between sequencing depth and the number of called ASVs in the ZymoBIOMICS mock community for each primer set, testing different minsize cutoff in the unoise3 algorithm for denoising**. Same data is shown in **A)** and **B)** however in **B)** the y-axis limit is set to ASV=45 to allow for better separation. For each sequencing platform, filled points (as shown in legend) indicate the number of ASVs with an exact match to a theoretical ASV, while open points represent the total number of ASVs. For each primer set, the dotted line indicates the number of theoretical ASVs based on curated Zmock reference sequences. Identical subsampling depths were applied across all datasets, and the final point represents the maximum number of available reads.

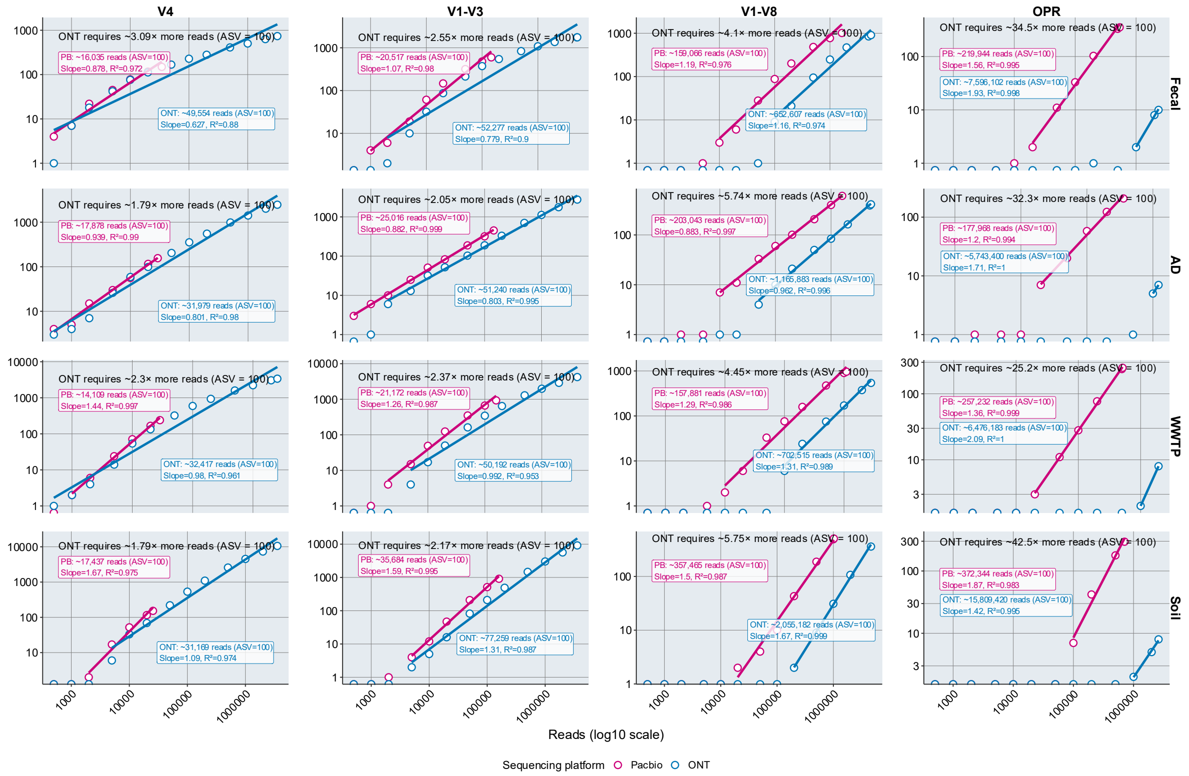


**Supplementary Fig. S7: ASV accumulation curves and linear-phase efficiency for PacBio and ONT across primer sets and sample types.** The number of recovered ASVs is plotted against sequencing depth on a log-log scale for PacBio and ONT. Linear-phase points were identified using primer-specific criteria, and linear models of log-transformed ASV counts versus log-transformed read depth were fitted to this region only (solid lines). For each panel, the fold-difference in reads required between ONT and PacBio (ASV = 100) is printed and for each sequencing platform the text box show the estimated read depth required to recover the ASV=100 and the slope and $R^2$ of the log-log linear model

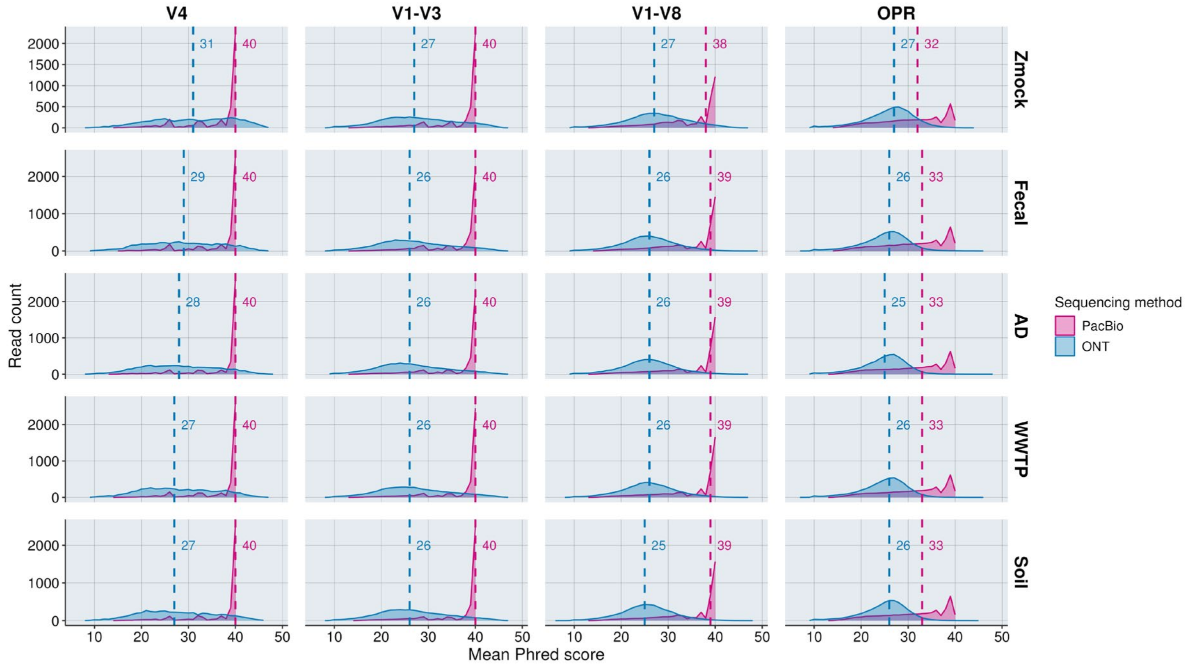


**Supplementary Fig. S8: Distribution of mean per-read Phred scores for PacBio and Oxford Nanopore (ONT) sequencing across all samples and primer sets.** For each dataset, reads were binned by their mean Phred score (bin size = 1) and plotted as area curves to visualize the quality distribution. All reads were primer trimmed before calculating Phred scores and a subsample of the trimmed reads were used with 10,000 reads for the V4 datasets and for the remaining datasets 100,000 reads and in these cases the y-axis values correspond to 10x read count. The mean per-read is shown separately for sample types (Zmock, Fecal, AD, WWTP, Soil) across the four primer sets (V4, V1-V3, V1-V8, OPR). Colors distinguish sequencing platforms. Dashed vertical lines indicate the weighted median Phred score with the values printed for each platform within each plot.

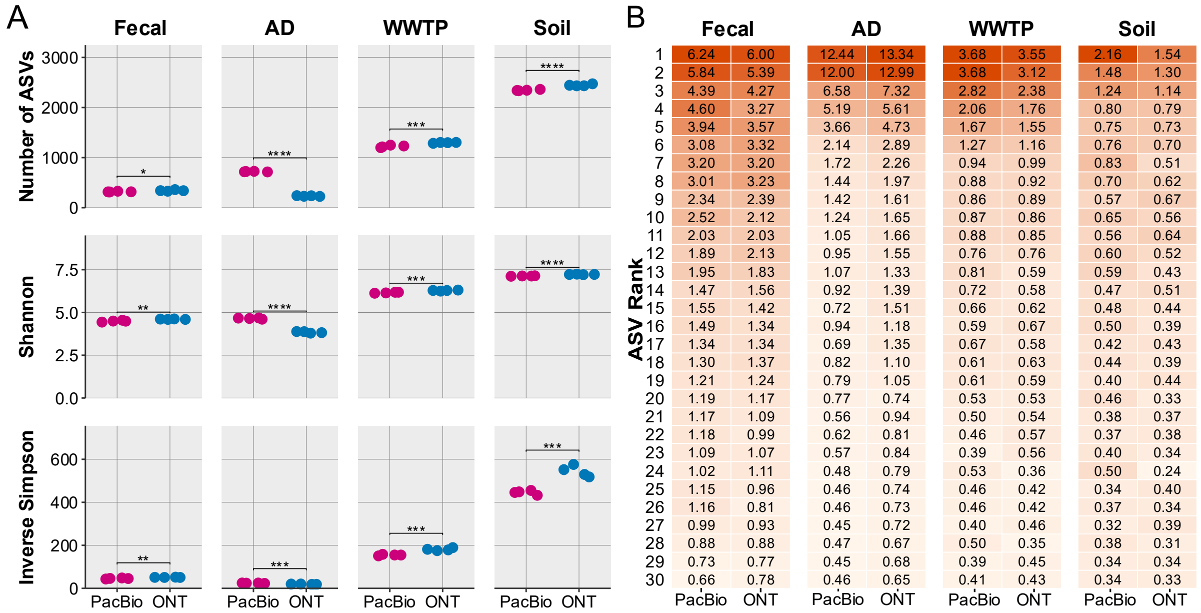


**Supplementary Fig. S9: Alpha-diversity and top 30 most abundant ASVs in the complex communities (V4 data). A)** Alpha-diversity indices; a t-test was performed to compare mean values between sequencing platforms (ns: not significant; *: p<0.05; **: p<0.01; ***: p<0.001). **B)** Relative abundance of the top 30 most abundant ASVs in each sample type with the relative abundance shown in percent. All analyses were performed on a dataset rarefied to 5,126 reads, corresponding to the sample with the fewest reads.

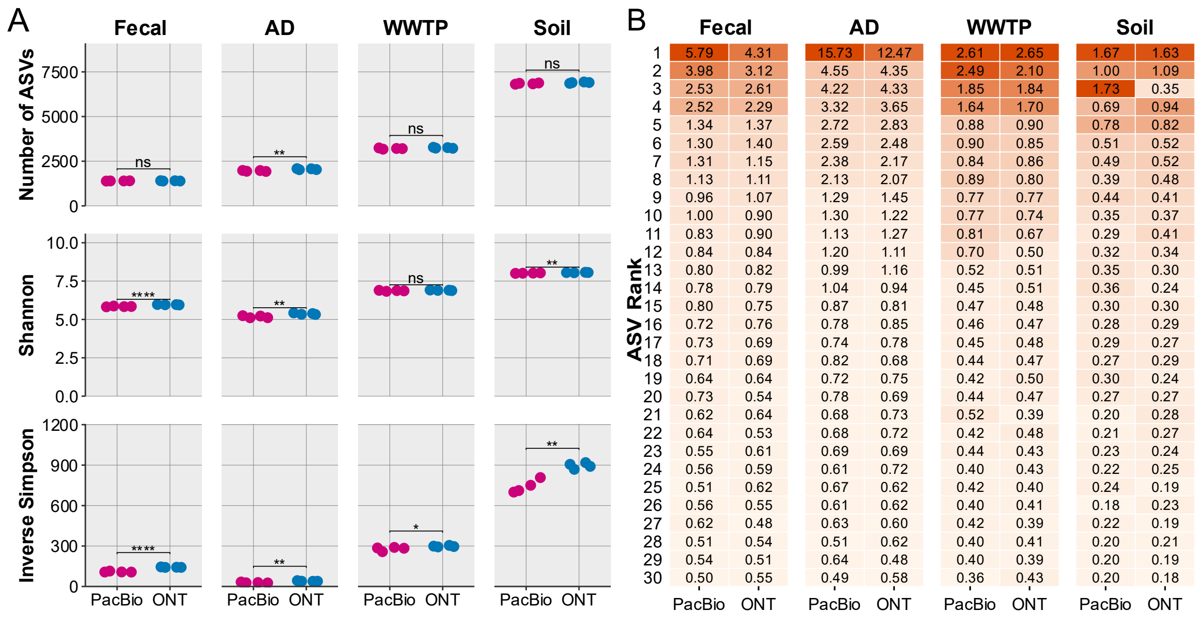


**Supplementary Fig. S10: Alpha-diversity and top 30 most abundant ASVs in the complex communities (V1-V3 data). A)** Alpha-diversity indices; a t-test was performed to compare mean values between sequencing platforms (ns: not significant; *: p<0.05; **: p<0.01; ***: p<0.001). **B)** Relative abundance of the top 30 most abundant ASVs in each sample type with the relative abundance shown in percent. All analyses were performed on a dataset rarefied to 29,083 reads, corresponding to the sample with the fewest reads.

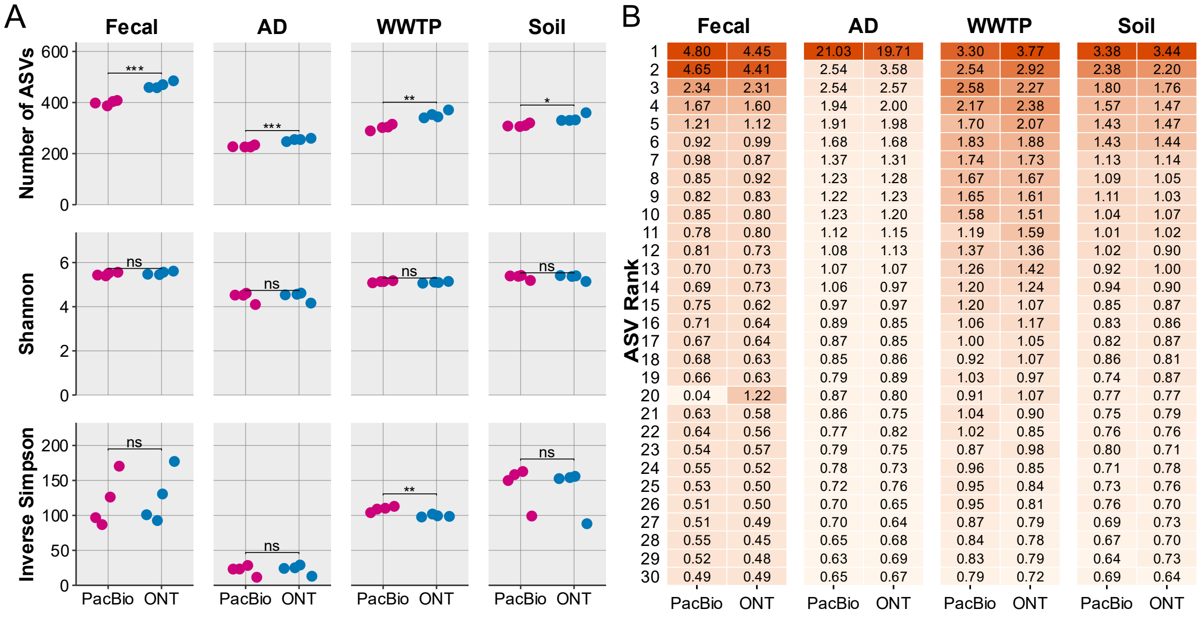


**Supplementary Fig. S11: Alpha-diversity and top 30 most abundant ASVs in the complex communities (OPR data). A)** Alpha-diversity indices; a t-test was performed to compare mean values between sequencing platforms (ns: not significant; *: p<0.05; **: p<0.01; ***: p<0.001). **B)** Relative abundance of the top 30 most abundant ASVs in each sample type with the relative abundance shown in percent. All analyses were performed on a dataset rarefied to 27,135 reads, corresponding to the sample with the fewest reads.

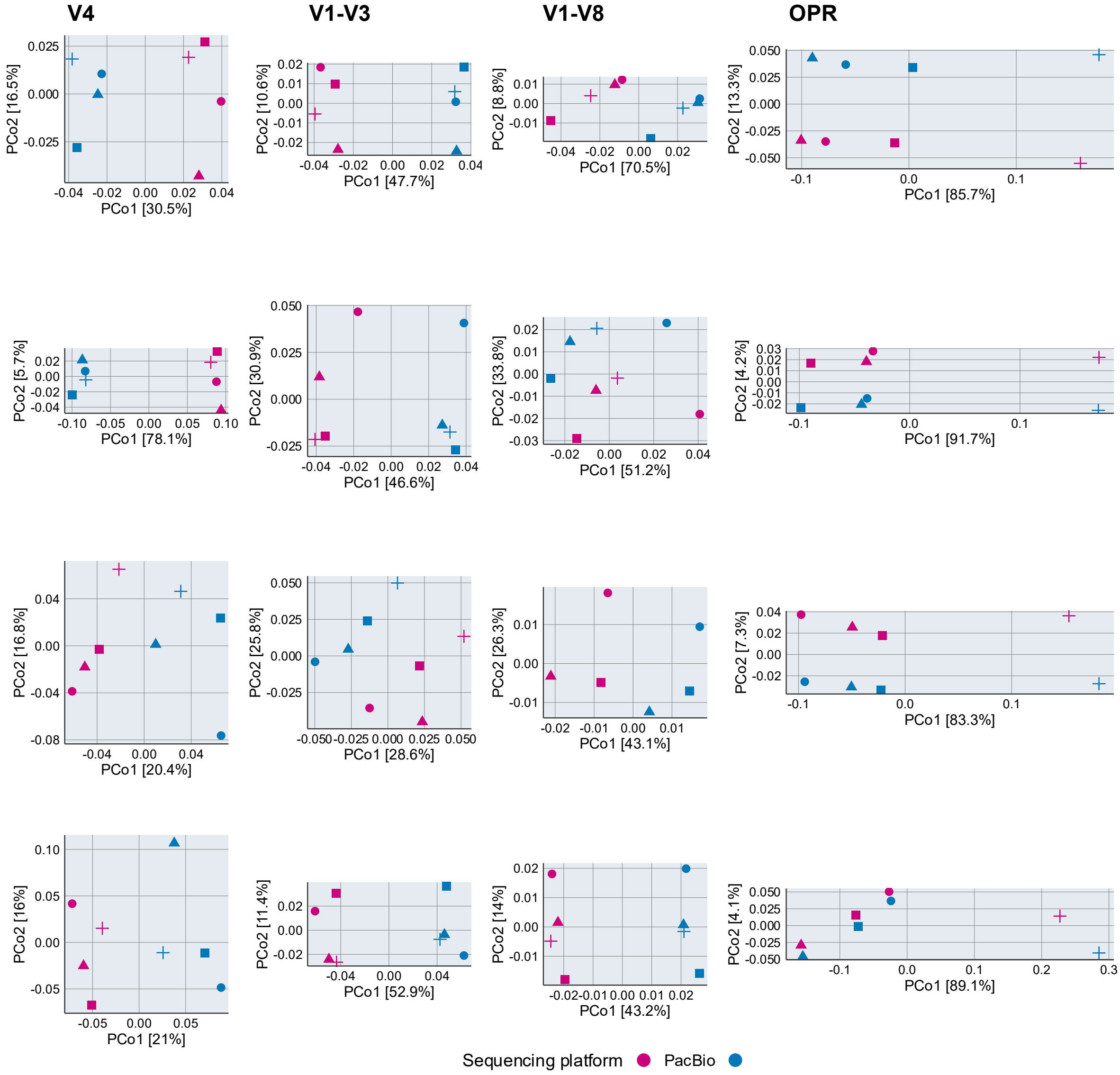


**Supplementary Fig. S12: Beta-diversity patterns for each primer set and sample type based on Bray–Curtis dissimilarities.** Shapes indicate biological replicates and colors denote sequencing platform. Analyses were performed on datasets rarefied to the sample with the fewest reads within each primer set (V4: 5,126–6,896 reads; V1–V3: 29,083–31,342 reads; V1–V8: 88,500–324,231 reads; OPR: 27,135–68,587 reads). Only ASVs with a relative abundance ≥ 0.1% were included.

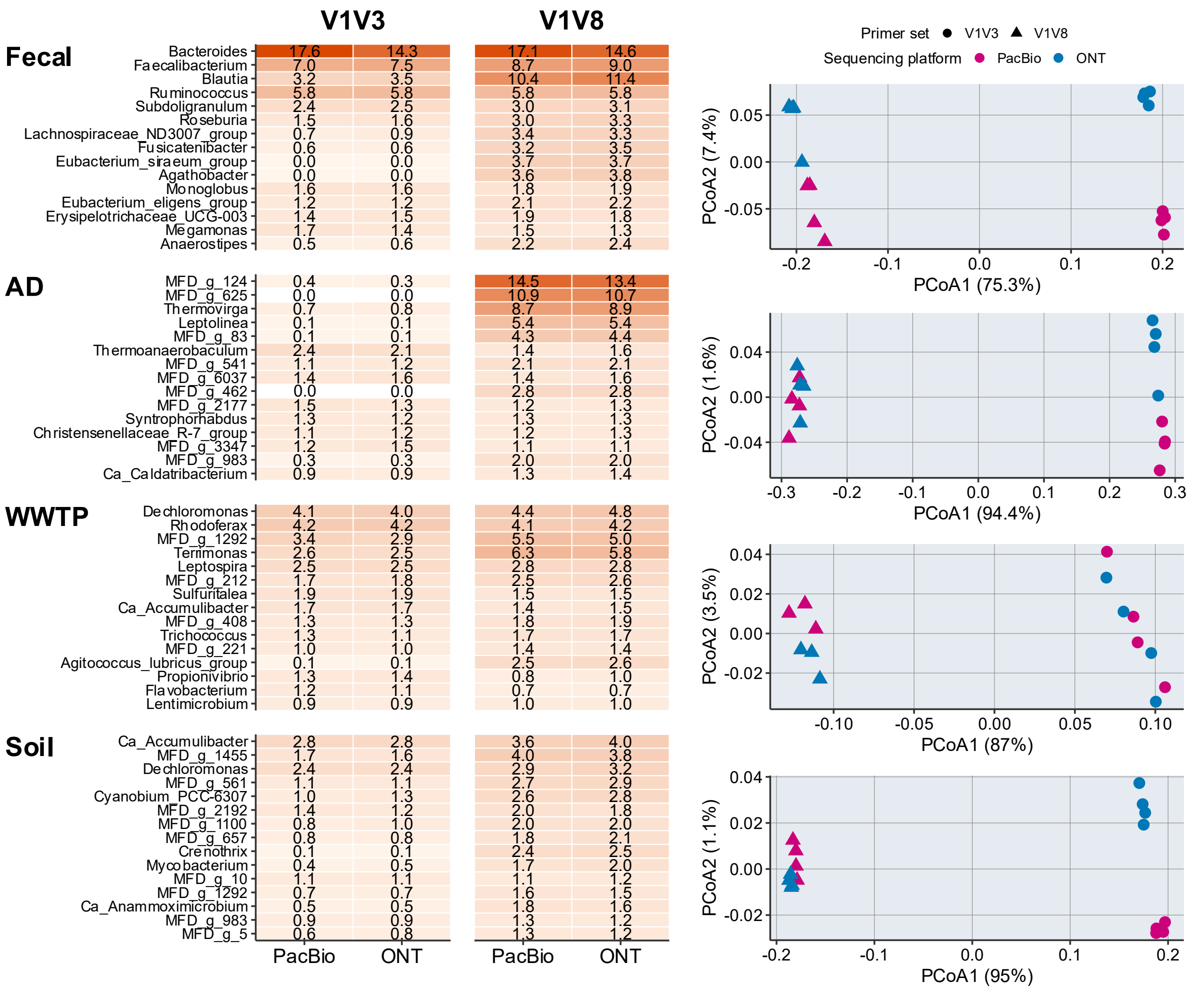


**Supplementary Fig. S13: Taxonomic diversity across V1-V3 and V1-V8 primers.** All analyses were performed on rarefied datasets with read counts matched to the sample with the fewest reads: 29083. **A)** Top 15 most abundant genera across each sample content. **B)** Beta diversity of samples at the genus level for each sample type based on Bray-Curtis distance matrices, including only ASVs classified to the genus level and with a relative abundance ≥ 0.1%. PERMANOVA results were assessed for the effects of sequencing platform (SP) and primer set (PS). Fecal: SP: $R^2$=0.07, $P$<0.246, PS: $R^2$=0.95, $P$<0.001; AD: SP: $R^2$=0.01, $P$<0.615, PS: $R^2$=0.97, $P$<0.001; WWTP: SP: $R^2$=0.02, $P$<0.62, PS: $R^2$=0.86, $P$<0.001; Soil: SP: $R^2$=0.01, $P$<0.638, PS: $R^2$=0.95, $P$<0.001.

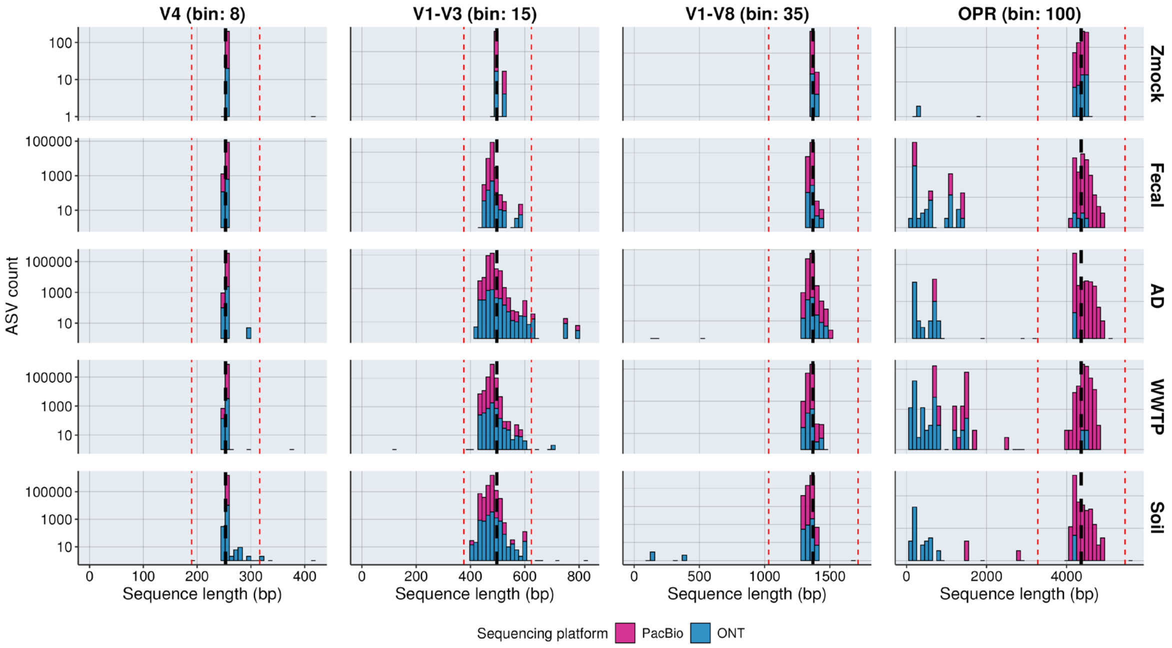


**Supplementary Fig. S14**: **Length distribution of amplicon sequence variants (ASVs) obtained from PacBio and Oxford Nanopore (ONT) sequencing across primer sets and five communities.** Histograms show the observed ASV sequence lengths for V4, V1-V3, V1-V8, and OPR across Zmock, Fecal, AD, WWTP, and soil communities, with PacBio (pink) and ONT (blue) stacked. Dashed red lines indicate the accepted amplicon length range (±25% of the median Zmock length), with the median itself marked by a black stippled line. Limits (minimum, median, maximum) for each primer set are (bp): V4 (190; 253; 316), V1-V3 (376; 497; 624), V1-V8 (1031; 1370; 1716), and OPR (3281; 4364; 5463). Histogram bin widths (bin) were adjusted for each primer set to account for differences in amplicon length, allowing sufficient resolution for longer amplicons.
